# Rod pathway blockade improves visual function in a mouse model of photoreceptor degeneration

**DOI:** 10.64898/2026.07.17.739110

**Authors:** Manoj M. Kulkarni, Jacqueline Gayet, Kristel Cosio, Amy Zheng, Teresa Puthussery, W. Rowland Taylor

## Abstract

Retinitis pigmentosa (RP) refers to a group of inherited photoreceptor degenerations characterized by the initial death of rods and secondary loss of cones. In animal models of RP, the ganglion cells (GCs) exhibit increased spontaneous activity that can mask residual photoreceptor signals and reduce the efficacy of treatments for vision restoration. The A2 amacrine cells (ACs) are inhibitory interneurons that propagate aberrant activity to GCs, but it remains unclear whether rod or cone pathways drive altered A2-AC signaling or whether the relative contributions of these pathways change with disease progression. We examined the circuit mechanisms underlying aberrant activity in the *rd10* mouse model and found that blocking rod pathway input to A2-ACs suppressed aberrant activity and improved the signal-to-noise ratio of residual cone-driven responses in On-α GCs. The results also indicate that the On- but not the Off-pathway exhibits circuit-level compensation during the loss of photoreceptor input. The results suggest that pharmacological blockade of the rod pathway may improve residual cone-mediated function and improve the efficacy of vision restoration in rod-cone photoreceptor degenerations.

## Introduction

Retinitis pigmentosa (RP) is a group of genetically heterogeneous inherited retinal degenerations characterized by progressive visual impairment and eventual blindness. Although rod photoreceptors harbor the pathogenic mutations causing rod degeneration and loss of night vision, the debilitating loss of high acuity daytime vision results from the subsequent death of cones. Since cone degeneration is relatively slow, and function can persist even in partially degenerate cones (Ellis et al. 2023; Scalabrino et al. 2022), therapeutic strategies that enhance the transmission of residual cone signals would be beneficial.

Photoreceptor death leads to a variety of downstream changes in the neural circuits of the inner retina. In rodent models of RP, the ganglion cells (GCs) exhibit aberrant activity that manifests as rhythmic oscillations or increased spontaneous firing (Zeck 2016; Euler and Schubert 2015; Stasheff 2008; Xiang et al. 2018; Trenholm and Awatramani 2015). Since this aberrant activity is uncorrelated with visual inputs, it can reduce the signal-to-noise ratio of residual cone-driven signals (Ivanova et al. 2016; Eleftheriou et al. 2017; Stasheff 2018; Toychiev et al. 2013). Similarly, aberrant activity will reduce the efficacy of therapies that restore light-sensitivity to the retina, for example with electrical prostheses or optogenetics (for review see: (Van Gelder et al. 2022; Baker and Flannery 2018; Lindner et al. 2022)). Thus, the saliency of residual cone-driven signals and the efficacy of vision restoration therapies could be enhanced by suppressing aberrant activity.

Studies in rodent models of retinitis pigmentosa (RP) indicate that A2-ACs play a key role in generating the aberrant activity that is seen in GCs (Margolis et al. 2014; Poria and Dhingra 2015). A2-ACs are inhibitory interneurons that play a central role in both scotopic and photopic visual processing (Demb and Singer 2012). Aberrant activity in A2-ACs has been proposed to arise from cell-intrinsic mechanisms or from gap junction connections between A2-ACs and ON-cone bipolar cells (CBCs) (Borowska, Trenholm, and Awatramani 2011; Choi et al. 2014; Ivanova et al. 2016; Eleftheriou et al. 2017; Stasheff 2018; Biswas et al. 2014; Menzler and Zeck 2011; Menzler, Channappa, and Zeck 2014). However, A2-ACs also receive direct excitatory input from the rod pathway via rod bipolar cells (RBCs), which undergo molecular, structural, and functional changes after rods degenerate (for review see (D’Orazi, Suzuki, and Wong 2014)). For example, RBCs lose dendritic mGluR6 receptors, remodel their dendrites to form ectopic synapses with cones, and may be involved in outer retinal oscillatory activity (Gargini et al. 2007; Puthussery et al. 2009; Strettoi et al. 2002; Haq et al. 2014; Fransen et al. 2015). Despite these changes, the extent to which RBCs drive aberrant activity in the degenerating retina remains unclear. Moreover, since rod death precedes cone death in RP, the contribution of the rod and cone pathways could change at different stages of degeneration.

Prior studies have shown that calcium-permeable AMPA receptors (CP-AMPARs) mediate transmission from RBC’s to AII AC’s in a variety of mammalian species, including primates (Mørkve, Veruki, and Hartveit 2002; Singer and Diamond 2003; R. S. Jones et al. 2014; Percival et al. 2022). Here, we use this pharmacology to evaluate the contribution of the rod pathway to aberrant activity in the *rd10* mouse, a model of human retinitis pigmentosa (Chang et al. 2002). We show that blocking excitatory input from RBCs to A2-ACs suppresses aberrant activity in downstream On-α GCs, thus improving the signal-to-noise ratio of residual cone-driven visual signals. The results also indicate the presence of pathway-selective circuit level compensation during degeneration.

## Results

### Scotopic input to A2-ACs is mediated by CP-AMPARs

Our first goal was to determine the contributions of the rod and cone pathways to aberrant A2-AC activity. To do so, we needed to selectively block signal input from the rod pathway, while preserving cone-driven inputs. Prior studies using paired recordings and agonist application have shown that CP-AMPARs mediate synaptic transmission from rod bipolar cells (RBCs) to A2-ACs (Mørkve, Veruki, and Hartveit 2002; Singer and Diamond 2003; Percival et al. 2022; Veruki, Mørkve, and Hartveit 2003), but the contribution of CP-AMPARs at this synapse under physiological conditions is less clear. Thus, we first tested whether we could block rod-driven input to A2-ACs in adult (P52-P68) wild-type (*wt*) mouse retinas using the CP-AMPAR antagonist, IEM-1460 (Magazanik et al. 1997). To do so, we measured flash-evoked excitatory postsynaptic currents (EPSCs) from A2-ACs in retinal slices maintained under dark-adapted conditions (Fig. 1A). IEM (50 μM) reversibly suppressed these EPSCs by 81% (Fig. 1B-D, mean ± s.d.: Ctrl, −170 ± 109 vs IEM, −32 ± 20 pA, Wilcoxon signed-rank test, *p* = 9.8×10^−4^, n = 11 cells) indicating that CP-AMPARs mediate most of the transmission from RBCs to A2-ACs under scotopic conditions. Thus, we used IEM to assess the role of the rod pathway in driving aberrant activity in the degenerating retina in subsequent experiments.

**Figure 1:**
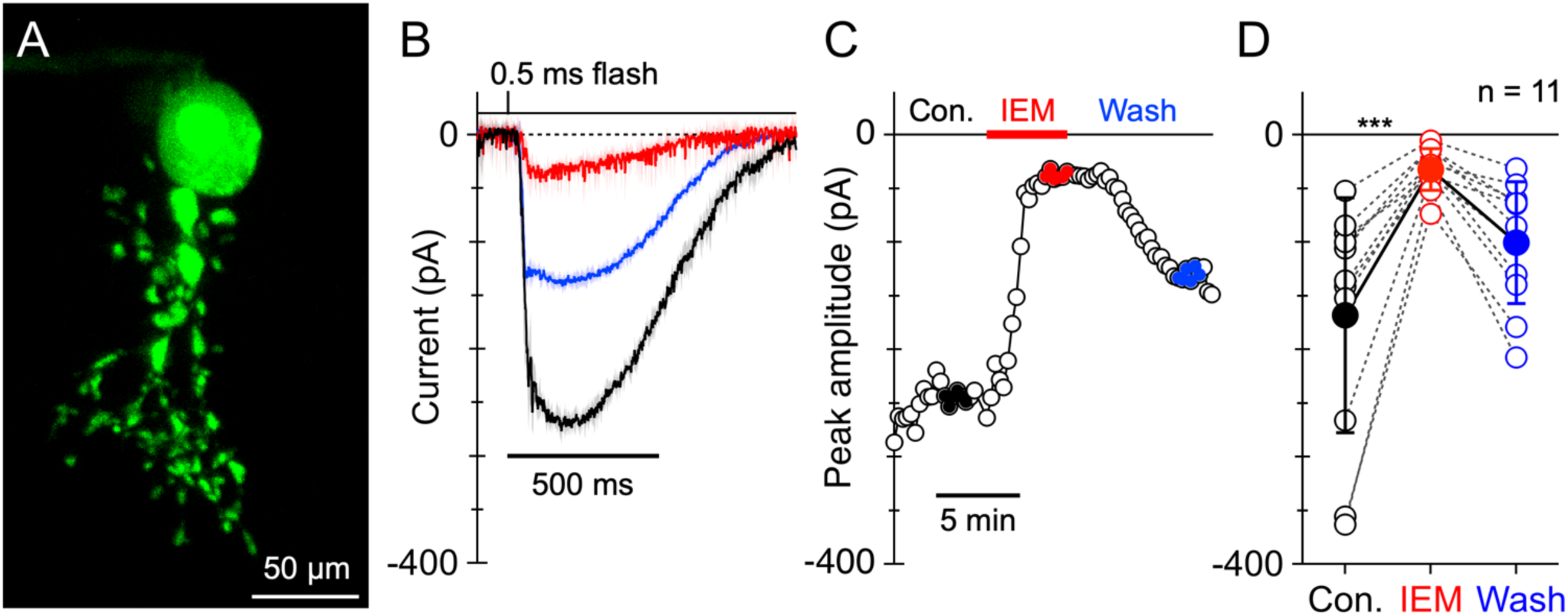
Scotopic input to A2-ACs is mediated by CP-AMPARs. **A.** Morphology of an A2-AC recorded from a slice preparation. **B.** Average EPSCs recoded at −70 mV (6 trials, solid symbols C) in an A2-AC in response to a 1.2 ms saturating light flash before (Control), during (IEM 50µM) and after washout (Wash) of IEM. Shading shows standard deviation. **C.** Peak amplitude of the light-evoked EPSCs as a function of time during application of IEM-1460 (50 µM). **D.** Peak amplitudes from 11 cells from 5 retinas. The filled symbols show mean ± s.d.. p = 9.8 x 10^−4^, Wilcoxon signed-rank test.

### The rod pathway depolarizes A2-ACs and contributes to aberrant activity in rd10 retina

A2-ACs contribute to aberrant activity in GCs in the *rd1* mouse model, however, a limitation of this model is that photoreceptor loss overlaps with the peak period of retinal circuit development (Trenholm et al. 2020; Choi et al. 2014; Margolis et al. 2014). In the *rd10* mouse, degeneration is relatively delayed and occurs more slowly after synapse development is complete, removing the confound of developmental overlap. The role of A2-ACs in driving aberrant activity in the *rd10* model has only been indirectly tested (Ivanova, Yee, and Sagdullaev 2015; Biswas et al. 2014; Menzler, Channappa, and Zeck 2014; Toychiev et al. 2013). Here, we used 50 µM IEM to test whether CP-AMPARs drive aberrant activity in A2-ACs in *rd10* retinas. We recorded excitatory postsynaptic potentials (EPSPs) from A2-ACs in “young” animals (defined as P21-25) before rod degeneration, and in “old” animals (defined as P35-150) after the peak period of rod degeneration (Gargini et al. 2007; Puthussery et al. 2009). We found a variety of activity patterns across different cells, some examples of which are illustrated in Fig. 2A. Analysis of the temporal power spectra of the membrane potential revealed peaks in many cells at frequencies between 4 to 10 Hz (Fig. 2B,C). IEM produced variable effects on the power spectra, including strong suppression (Fig. 2B, upper), a shift to lower frequencies (Fig. 2B, middle), or minimal effect (Fig. 2B, lower). IEM suppressed oscillatory activity more markedly in young versus old animals (Fig. 2C) but did not eliminate the oscillatory peaks at either time. IEM also hyperpolarized A2-ACs by 10.7 mV in young animals (Fig. 2D, control −56.6 ± 2.2 vs IEM −67.3 ± 11.3 mV, Wilcoxon signed-rank test, *p* = 0.0034, n = 12 cells) and 6.4 mV in old mice (Fig. 2D, control −57.6 ± 2.4 vs IEM −64.0 ± 6.4 mV, Wilcoxon signed-rank test, *p* = 2.44 x 10^−4^, n = 13 cells), indicating that a tonic excitatory drive from RBCs to A2-ACs persists in the *rd10* retina after rods have degenerated.

**Figure 2:**
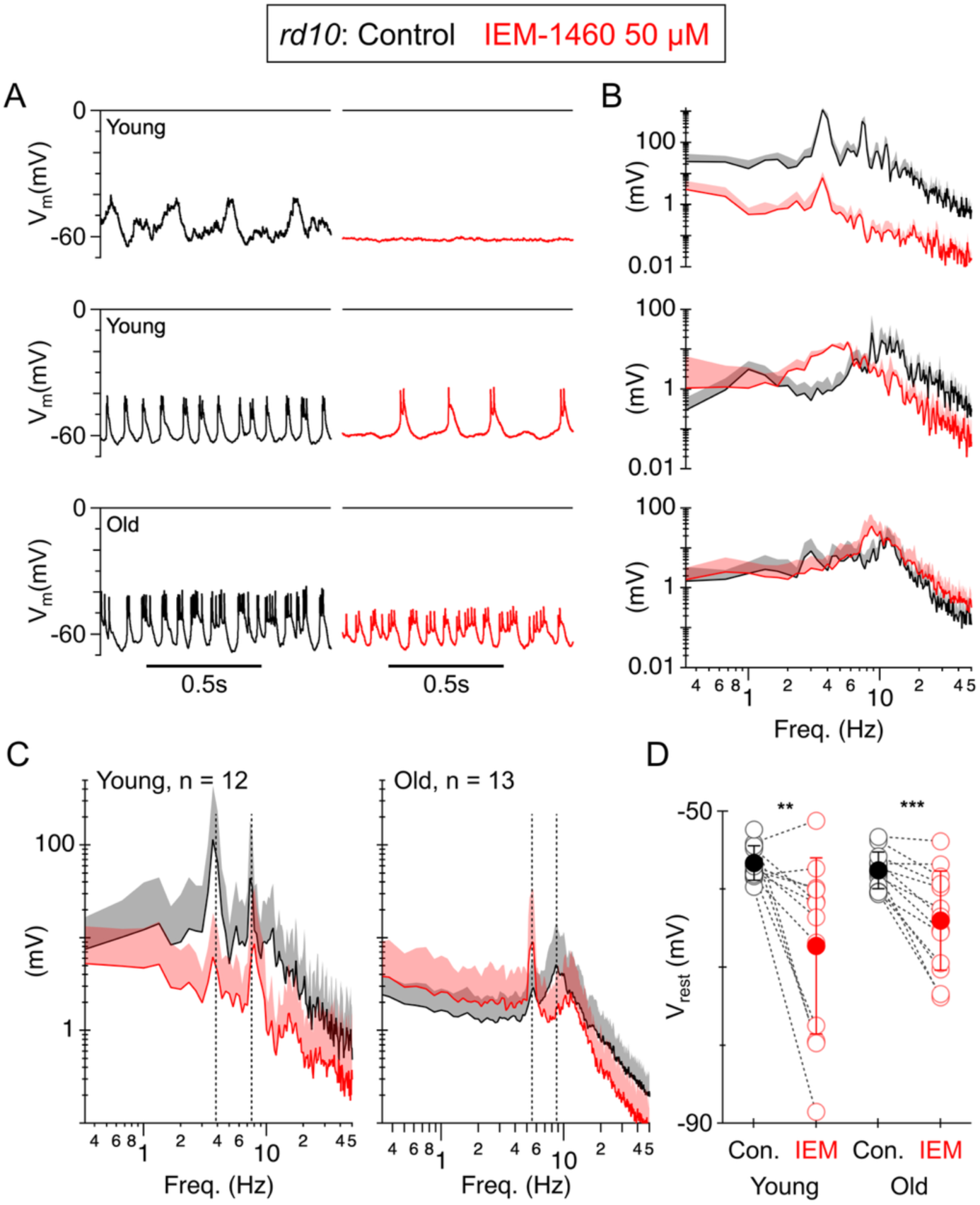
The rod pathway depolarizes A2-ACs and contributes to aberrant activity in the *rd10* retina. **A.** Three example A2-ACs cells from *rd10* retina showing spontaneous voltage oscillations in control (black) and in the presence of 50 µM IEM (red). **B.** Power spectral density (PSD) estimates were obtained by averaging temporal power spectra from five 3 second segments of voltage recording. **C.** Average PSDs from A2-ACs in young and old animals. **D.** Mean resting potential for the same cells shown in *C.* Solid symbols show mean open symbols show individual cells. Statistical comparisons in D were made using Wilcoxon signed-rank test. Shading and error bars show ±1 s.d..

### Increased baseline firing in the On pathway in rd10 vs wild-type retina

Aberrant activity in A2-ACs is thought to propagate to GCs by signaling through intervening On and Off cone bipolar cells (Margolis et al. 2014; Choi et al. 2014; Borowska, Trenholm, and Awatramani 2011). To test the impact of degeneration on downstream GCs, we focused on three GC types: On-α sustained (On), Off-α sustained (sOff), and Off-α transient (tOff). The α-type GCs are the likely orthologs of primate midget (sustained α) and parasol (transient α) GCs (Hahn et al. 2023). We targeted α-type GCs based on their large soma size (Murphy and Rieke 2006; M. B. Manookin et al. 2008; van Wyk, Wässle, and Taylor 2009; Grimes et al. 2022).

Consistent with prior studies (Ivanova et al. 2016; Stasheff, Shankar, and Andrews 2011; Xiang et al. 2018), baseline spike rates in On-α GCs, measured during 3 s immediately prior to the stimulus, were higher in *rd10* retinas than age-matched *wt* controls (Fig. 3A,B). In young *wt* retinas the baseline firing rate averaged 5.8 Hz and increased by 136% to 13.6 Hz in the age-matched *rd10* group. Similarly, in old *wt* retinas, the average baseline firing rate average 10.3 Hz and increased by 130% to 24.1 Hz in the age-matched *rd10* group (Fig. 3C,D, Table 1). In contrast, baseline spike rates in Off-α GCs were not significantly different in either strain at either age (Fig. 3A-D, Table 1).

**Figure 3:**
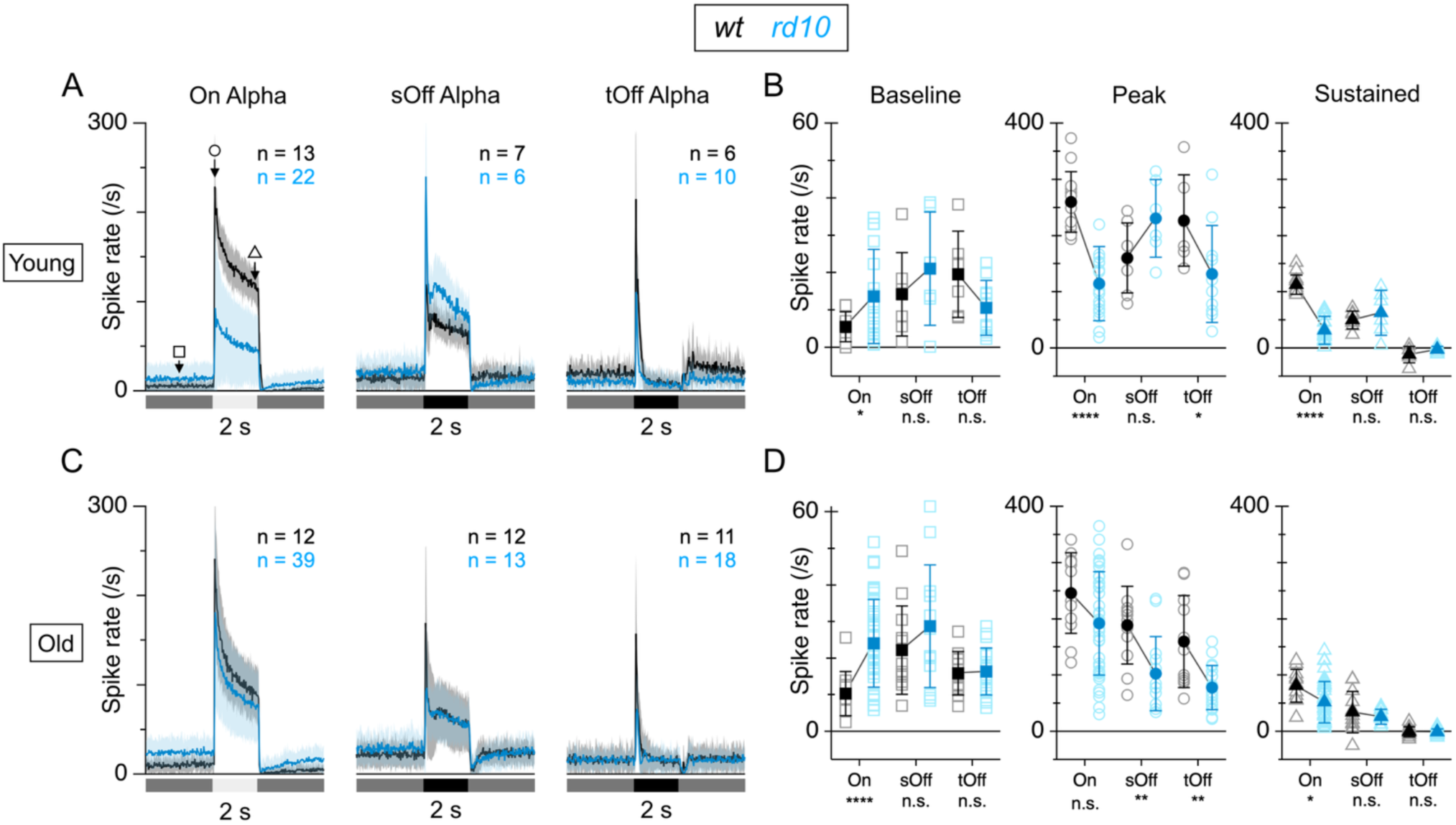
Increased baseline firing in ON-α GCs in *rd10* vs *wt* retina. Extracellular recordings of baseline and light-evoked firing in α GCs from young (**A,B**) and old (**C-D**) mice in *wt* control groups (black) and age-matched *rd10* groups (blue). Full-field stimuli, 100% contrast. **A.** Average peri-stimulus spike-time histograms (PSTHs) in α GCs recorded in young mice. The number of cells for the averages are shown in each panel. Bars below show timing and contrast of spot stimuli centered on the receptive field. **B.** Summary data for the baseline, peak, and sustained firing rates for the three GC types. The measurement time-points for each are shown by the corresponding symbols on the On-α PSTHs in *A*. The peak and sustained amplitudes are baseline subtracted. Solid symbols show mean ± s.d., open symbols show individual cells. Statistical comparisons by two-tailed t-test. **C,D.** Same format as **A,B** for the “old” mouse cohorts. Error bars and shading show ± 1 s.d..

**Table 1:** Comparisons across genotype and age for spike rates (Hz) recorded in *wt* and *rd10* GCs in young and old animals (Fig. 3). See Methods for definitions of baseline, peak, and sustained. The p-values were calculated from unpaired T-tests using Welch’s correction. Negative spike-rates indicate a suppression of firing relative to the initial pre-stimulus baseline level. Bold text highlights significant changes (p < 0.05).

|  | Young |  | Old |  | *wt* vs *rd10* : p, %change |  | Young vs Old : p, %change |  |
| --- | --- | --- | --- | --- | --- | --- | --- | --- |
|  | *wt* | *rd10* | *wt* | *rd10* | Young | Old | *wt* | *rd10* |
| On-α | | | | | | | | |
| Baseline | 5.8 ± 4.1<br>(n=12) | 13.6 ± 12.6<br>(n=22) | 10.3 ± 6.0<br>(n=12) | 24.1 ± 12.0<br>(n=39) | **0.012, 140%** | **4.9x10<sup>-6</sup>, 130%** | **0.044, 78%** | **0.0029, 77%** |
| Peak | 236 ± 42 | 115 ± 66 | 246 ± 72 | 192 ± 92 | **2.3x10<sup>-7</sup>, -51%** | 0.066, -22% | 0.68, 4% | **3.5x10<sup>-4</sup>, 68%** |
| Sustained | 113 ± 18 | 32.1 ± 25.0 | 81.0 ± 28.9 | 52.0 ± 36.6 | **9.6x10<sup>-12</sup>, -72%** | **0.016, -36%** | **0.0044, -28%** | **0.015, 62%** |
| sOff-α | | | | | | | | |
| Baseline | 14.3 ± 11.1<br>(n=7) | 21.1 ± 15.2<br>(n=6) | 22.2 ± 12.1<br>(n=12) | 28.7 ± 16.8<br>(n=13) | 0.38, 48% | 0.28, 29% | 0.17, 56% | 0.35, 36% |
| Peak | 160 ± 62 | 231 ± 69 | 189 ± 69 | 103 ± 66 | 0.083, 44% | **0.0044, -46%** | 0.38, 18% | **0.0038, -55%** |
| Sustained | 50.1 ± 15.9 | 74.3 ± 31.8 | 34.3 ± 36.8 | 25.9 ± 13.5 | 0.17, 48% | 0.47, -24% | 0.21, -32% | 0.075, -59% |
| tOff-α | | | | | | | | |
| Baseline | 19.6 ± 11.5<br>(n=6) | 10.6 ± 7.3<br>(n=10) | 15.9 ± 5.9<br>(n=11) | 16.3 ± 6.5<br>(n=18) | 0.13, -46% | 0.87, 3% | 0.50, -19% | 0.054, 54% |
| Peak | 227 ± 81 | 132 ± 86 | 160 ± 82 | 77.9 ± 39.2 | **0.048, -42%** | **0.0011, -51%** | 0.13, -30% | 0.086, -41% |
| Sustained | -11.4 ± 14.6 | -2.7 ± 5.4 | -0.4 ± 8.6 | -0.9 ± 5.9 | 0.21, -76% | 0.85, 150% | 0.13, -97% | 0.42, -67% |

### Degeneration affects light-evoked responses in On and Off pathways differently

We assessed light-evoked responses by stimulating GCs with 2 s full-field, 100% contrast light and dark flashes from a grey background (see Methods). As expected for extensive loss of rods and progressive loss of cones during degeneration, the peak light-evoked responses were reduced in the *rd10* retinas relative to age-matched *wt* retinas (Fig. 3). In the young *rd10* retina, peak light-evoked responses in On-α GCs averaged 115 Hz, which is 51% lower than the 236 Hz average response seen in age-matched *wt* controls (Fig. 3A,B, Table 1). The average peak light-evoked responses in young sOff-α GCs, were not significantly different in *rd10* and *wt* retinas (*rd10* 231 Hz, *wt* 160 Hz) but were 42% smaller in tOff-α GCs (*rd10* 132 Hz, *wt* 227 Hz. Fig. 3A-B, Table 1). In the old *rd10* retina, the peak responses for the On-α GCs were not significantly different from *wt* controls (*rd10* 246 Hz, *wt* 192 Hz), unlike the Off-α GCs which were uniformly smaller; old sOff-α GCs and tOff-α GCs were 46% (*rd10* 103 Hz, *wt* 189 Hz) and 51% (*rd10* 78 Hz, *wt* 160 Hz) smaller than the *wt* controls (Fig. 3C-D, Table 1). The apparent lack of effect of degeneration on the baseline spiking in the Off-α GCs, compared with the On-α GCs suggests that degeneration affects the two pathways differently.

To further investigate this difference, we analyzed the data in Fig. 3 within each phenotype at the two ages. In the *wt* and *rd10* groups, baseline spiking of the On-α GCs increased similarly with age; by 78% in the *wt* group and 77% in the *rd10* group (Table 1). Baseline firing in the Off-pathway cells didn’t change significantly with age in either the *wt* or *rd10* groups. Peak light-evoked responses were stable with age in the *wt* retinas, while the sustained component became 28% smaller. In contrast light-evoked responses actually increased with age in *rd10* retinas; the peak response increased by 68% and the sustained component by 62%. Notably, light-evoked responses of the Off-α GCs the *wt* and *rd10* groups generally decreased with age (Table 1). This selective increase in the On-pathway with age points to adaptive plasticity that is not evident in the Off-pathway.

### Blocking the rod pathway suppresses baseline firing in ON-α GCs in the rd10 retina

The results predict that blocking rod pathway input from RBCs to A2-ACs should suppress aberrant activity in downstream GCs. We tested this prediction by applying 50 µM IEM and measuring the baseline spiking and light-evoked responses in the α-GCs in the *rd10* retina (Fig. 4). In On-α GCs IEM suppressed baseline spiking by 86% (control 19.7 Hz, IEM 2.7 Hz) and 71% (control 24.2 Hz, IEM 7.0 Hz) in young and old retinas respectively (Table 3). IEM had no effect on baseline spiking in either Off-α GC type in young retinas, and had variable effects in old retinas, increasing baseline spiking by 52% (control 15.8 Hz, IEM 24.1 Hz) in sOff-α GCs and decreasing it by 23% (control 17.5 Hz, IEM 13.4 Hz) in tOff-α GCs. IEM did not alter the peak or sustained light-evoked responses in On-α GC or tOff-α GC types (Fig 4A-D), but suppressed peak responses in young (22%) (control 237 Hz, IEM 185 Hz) and old (25%) (control 72.5 Hz, IEM 54.4 Hz) sOff-α GCs. Overall, these results are consistent with the notion that CP-AMPA receptors don’t play a major role in CBC to GC signal transmission. The selective suppression of baseline spiking in the On-pathway with modest effects on the light-evoked responses, suggests that IEM could improve the signal-to-noise ratio (SNR) for the residual light-driven responses in *rd10* retinas.

**Figure 4:**
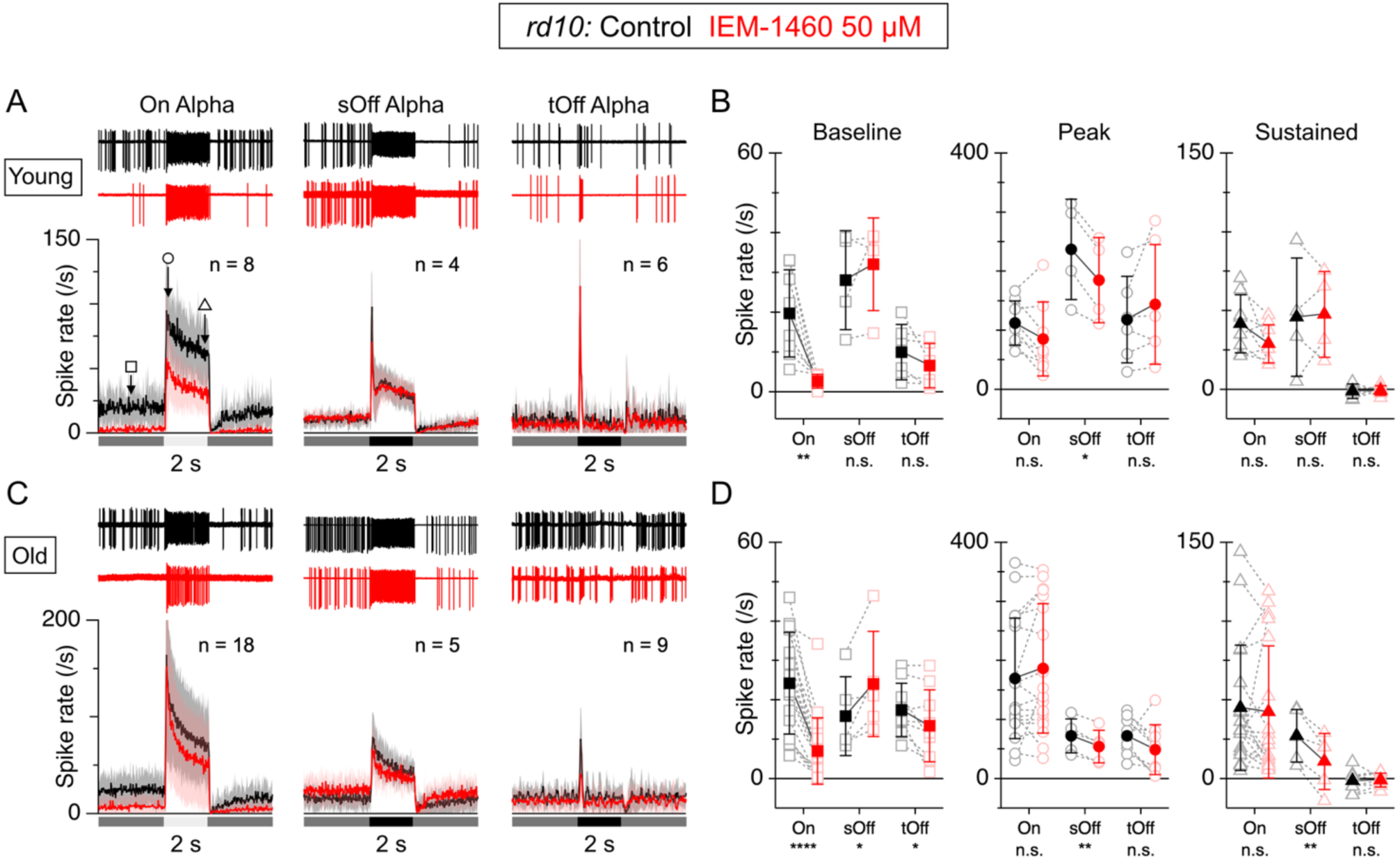
Rod pathway blockade hyperpolarizes A2-ACs and suppresses baseline firing in ON-GCs. Extracellular recordings of baseline and light-evoked firing in On-α GCs from young (**A,B**) and old (**C-D**) *rd10* mice in control conditions (black) and after IEM application (red). Full-field stimuli, 100% contrast. **A.** Average peri-stimulus spike-time histograms (PSTHs) in three α GC types recorded in young mice before and after IEM application. The number of cells for the averages are shown in each panel. Bars below show timing and contrast of a spot stimulus centered on the receptive field. **B.** Summary data showing the baseline, peak, and sustained firing rates for the three α GC types. The measurement time-points for each are shown by the corresponding symbols on the On-α GC PSTHs in **A**. The peak and sustained amplitudes are the changes in firing rate produced by the stimulus. Solid symbols show mean, open symbols show the individual cells. **C,D.** Same format as **A,B** for the “old” mouse cohorts. Error bars and shading show ± 1 s.d..

**Table 2:** Effect of glutamate receptor blocker, IEM (Fig. 4). Comparison of spike rates (Hz) recorded in *rd10* GCs, in control and during application of 50 µM IEM. See Methods for definitions of baseline, peak, and sustained. The *p*-values were calculated from paired T-tests. Negative spike-rates indicate a suppression of firing relative to the initial pre-stimulus baseline level. Bold text highlights significant changes (*p* < 0.05).

|  | Young |  | Old |  | Control vs IEM: p, %change |  |
| --- | --- | --- | --- | --- | --- | --- |
|  | Control | IEM | Control | IEM | Young | Old |
| On-α | | | | | | |
| Baseline | 19.7 ± 10.9 (n=8) | 2.7 ± 1.8 | 24.2 ± 12.9 (n=18) | 7.0 ± 8.4 | **0.0051** , -86% | **2.1x10<sup>-6</sup>** , -86% |
| Peak | 112 ± 37 | 85.6 ± 62.7 | 170 ± 102 | 187 ± 110 | 0.085 , -24% | **0.044** , 14% |
| Sustained | 41.6 ± 18.5 | 29.0 ± 12.0 | 45.0 ± 39.7 | 42.3 ± 42.0 | 0.078 , -30% | **0.037** , -33% |
| sOff-α | | | | | | |
| Baseline | 28.1 ± 12.5 (n=4) | 32.1 ± 11.6 | 15.8 ± 10.0 (n=5) | 24.1 ± 13.3 | 0.43 , -22% | **0.044** , 52% |
| Peak | 237 ± 85 | 185 ± 72 | 72.5 ± 28.9 | 54.4 ± 27.7 | **0.012** , 22% | **0.0020** , -25% |
| Sustained | 45.8 ± 37.6 | 47.6 ± 27.3 | 27.2 ± 16.8 | 10.8 ± 17.8 | 0.88 , -30% | **0.0076** , -60% |
| tOff-α | | | | | | |
| Baseline | 10.0 ± 7.0 (n=6) | 6.7 ± 5.5 | 17.5 ± 6.7 (n=9) | 13.4 ± 9.2 | 0.14 , -33% | **0.037** , -23% |
| Peak | 118 ± 73 | 144 ± 102 | 72.2 ± 31.0 | 49.2 ± 42.2 | 0.26 , 22% | 0.11 , -32% |
| Sustained | -1.1 ± 4.7 | -0.8 ± 3.0 | -1.4 ± 6.7 | -0.9 ± 4.3 | 0.92 , -27% | 0.75 , -38% |

### Rod pathway blockade improves signal-to-noise ratio in On-α GCs in rd10 retina

To test whether IEM could improve the SNR in On-α GCs in old *rd10* retina, we measured contrast-response functions before and after applying 5 μM IEM-1460 (Fig. 5). Although the higher concentration largely spared light-evoked activity, we used this lower concentration to ensure selective suppression of baseline firing. At 5 µM, IEM suppressed baseline firing by 81% (control 29.2 ± 25.9 Hz vs IEM 5.5 ± 4.5 Hz, n = 8, *p* = 0.0078, Wilcoxon signed rank, Fig. 5A,B), similar to the effect of the 50 µM dose (Fig 4). While 5 µM IEM did not affect peak firing at high contrasts (>50%) it significantly suppressed peak rates at contrasts between 5% and 50% (Fig. 5C, n = 8, *p* = 0.0056, cluster permutation analysis, see Methods for details). The sustained firing rate was unaffected by IEM application (Fig. 5F). IEM increased the half-maximal contrast (C_1/2_) for both the peak and sustained light response (Fig. 5D, peak: control 22.6 ± 6.4% vs IEM 33.7 ± 9.1%, *p* = 0.00137; Fig. 5G, sustained: control 20.8 ± 9.0% vs IEM 36.6 ± 12.5%, *p* = 0.00109). An increase in *C_half_* will correspond to an increase in the dynamic range, defined as the change in contrast, **Δ**C, between 10% and 90% of *R_max_* (**Δ**C_10-90_= 8.89\**C_half_*).

**Figure 5:**
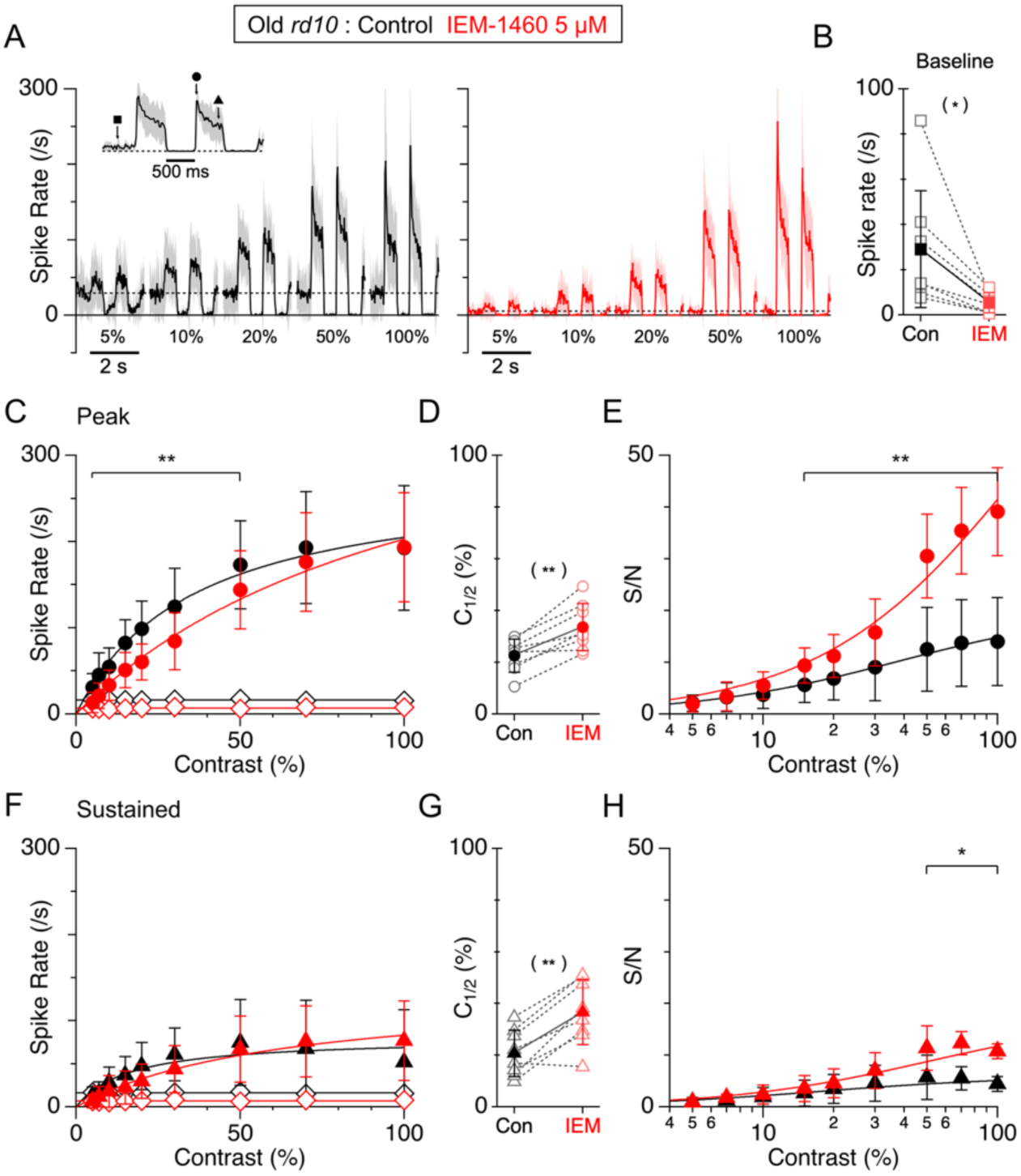
Rod pathway blockade increases SNR and shifts half-maximal contrast to higher levels in On-α GCs from *rd10* retina. Effect of IEM on contrast sensitivity of On-α GCs in old *rd10* retina. A. Average PSTHs (n=8 cells) in control (black) and in the presence of 5 µM IEM (red). The inset, showing the response at 100% contrast, illustrates the time-points for measuring the baseline, peak, and sustained amplitudes. The stimulus was a 200 µm diameter spot square-wave modulated at 1 Hz at the contrasts indicated beneath the traces. B. Summary data showing the effect of 5 µM IEM on the baseline spiking rate. Light symbols show the baseline firing, averaged across all contrasts, for each cell. C,F. Average response amplitudes as a function of stimulus contrast for the peak and sustained time-points. The open diamonds show the noise levels calculated as the standard deviation of the baseline firing rate preceding the light stimulus. D,G. Respective *C_1/2_* values (see Methods). Solid symbols show mean, open symbols show the individual cells. Statistical comparisons by paired t-test. E,H: SNR as a function of stimulus contrast for the corresponding amplitude measurements. Error bars and shading show ± 1 s.d..

A similar analysis of the SNR as a function of stimulus contrast found that IEM increased the SNR in a contrast-dependent manner for both the peak and sustained components of the light-response (Fig. 5E & H). The SNR for the peak response was increased at contrasts from 15 to 100% by an average of 2.2-fold (Fig. 5E, n = 8, *p* = 0.0084, cluster permutation analysis), while the SNR for the sustained component was increased at contrasts from 50 to 100% by an average of 2.8-fold (*p* = Fig. 5H, 0.0144, cluster permutation analysis). These results demonstrate that IEM can improve SNR in On-α GCs in old *rd10* retina.

An alternative approach to improve the SNR for residual responses might be to suppress inhibition and thereby potentiate the effects of excitatory transmission. We tested this idea by blocking glycinergic and GABAergic transmission using 1 μM strychnine, 5 μM GABAzine (GABA_A_ antagonist), and 100 μM TPMPA (GABA_C_ antagonist) in On-α GCs in old *rd10* retinas (Fig. 6). Unlike IEM, blocking inhibition had no effect on the baseline firing rate (control 20.5 ± 10.2 Hz vs inhibitory block 22.9 ± 7.7 Hz, *n = 4, P* = 0.46, paired T-test, Fig. 6A,B). As expected, the average peak light-evoked firing rate increased after blocking inhibition (Fig. 6C), however the effect was not significant overall (P = 0.062). The increase in the average peak SNR after inhibitory block (Fig. 6E) was also not significant (P = 0.087). Blocking inhibition had no effect on the sustained component of the light responses. In contrast to IEM, blocking inhibition did not produce a rightward shift in half-maximal contrast values (Fig. 6D).

**Figure 6:**
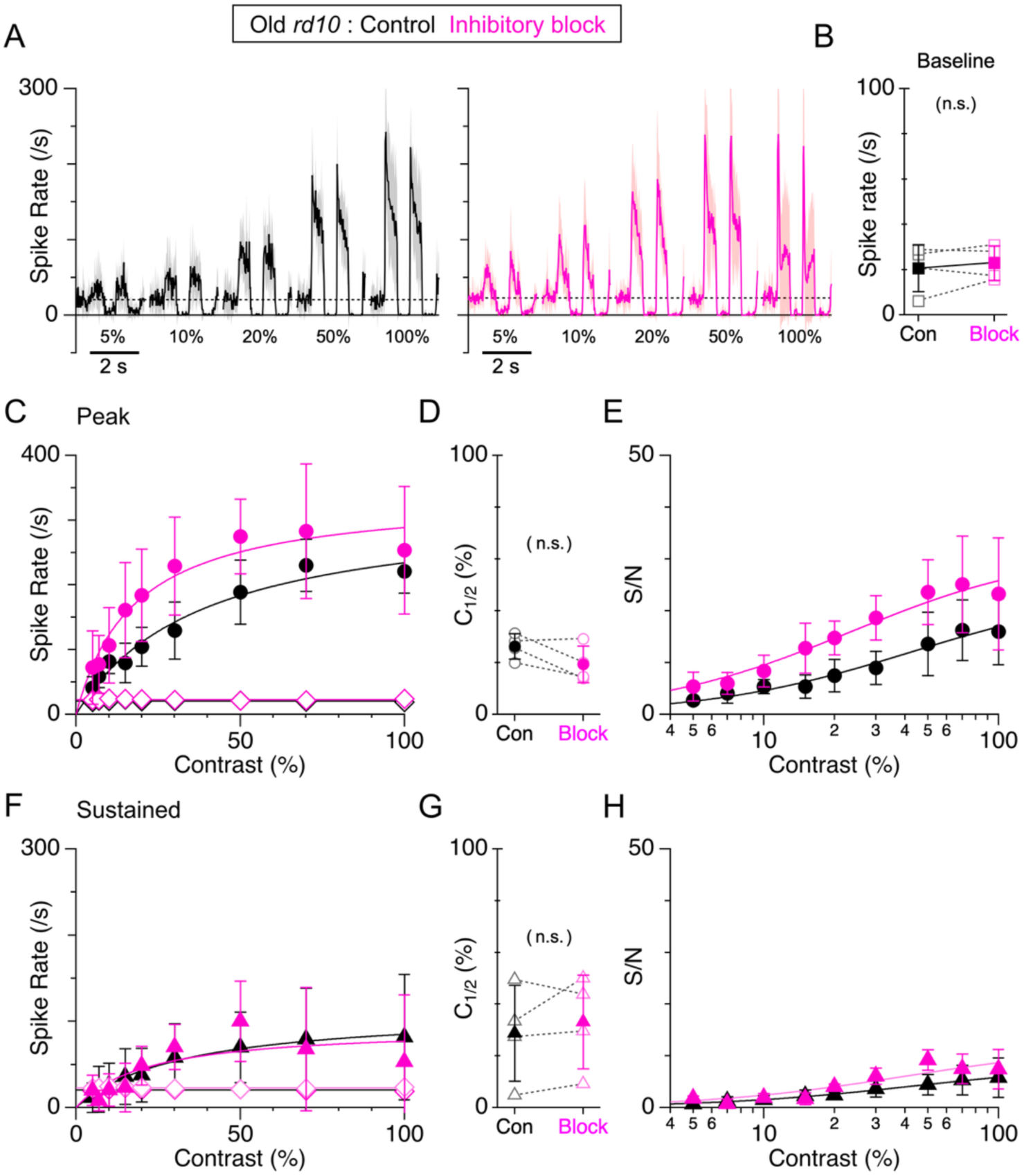
Inhibitory blockade increases SNR in On-α GCs in *rd10* retina. Effect of inhibitory blockade (GABA_A_, GABA_C_ and glycine receptors) on contrast sensitivity in On-α GCs from the old *rd10* retina. Average data from 4 cells. The format is the same as for Fig. 5. Data in the presence of inhibitory blockers is shown in magenta.

### CP-AMPARs are expressed in a subset of glycinergic amacrine cells, but not GCs

Normally CP-AMPARs are expressed by A2-ACs, A17 amacrine cells, and horizontal cells (Mørkve, Veruki, and Hartveit 2002; Singer and Diamond 2003; R. S. Jones et al. 2014; Chávez, Singer, and Diamond 2006; Osswald, Galan, and Bowie 2007; Cueva Vargas et al. 2015; Percival et al. 2022), while ON-α GCs show little or no expression (Wen et al. 2018; Sladek and Nawy 2020). However, CP-AMPAR expression can be upregulated in GCs during pathological conditions such as glaucoma, ocular hypertension, and ischemic injury (Cueva Vargas et al. 2015; Wen et al. 2018; Sladek and Nawy 2020; Wang, Carroll, and Nawy 2014). To control for possible up-regulation in *rd10* retinas, we used an established cobalt (Co^2+^) staining method to compare CP-AMPAR localization in “old” *wt* and *rd10* retinas. Co^2+^ passes through CP-AMPARs and accumulates inside cells where it can be precipitated and visualized (Aurousseau, Osswald, and Bowie 2012; Osswald, Galan, and Bowie 2007). We applied 5 mM Co^2+^ and stimulated uptake by applying 10 mM L-glutamate to activate glutamate receptors.

Many amacrine cell somata located at the border of the inner nuclear layer and inner plexiform layer (Fig. 7A-B) showed evidence of Co^2+^ uptake. Most cobalt-loaded cells expressed the glycine transporter, GlyT1, and had somatic and dendritic morphology characteristic of A2-ACs (Veruki and Hartveit 2002; Pérez de Sevilla Müller et al. 2017). Although A2-ACs can be identified with antibodies for the transcription factor, Prox1 (Fig. 7C), this nuclear protein was masked by the dense cobalt signal in L-Glu treated retinas. We confirmed that Co^2+^ loading occurs through AMPA receptors by showing that co-application of the AMPAR antagonist GYKI-53655 (80 μM) completely suppressed Glu-dependent Co^2+^ loading. The density and soma position of A2-ACs is evident with Prox1+ staining in samples treated with the AMPAR antagonist. These results confirm prior reports of CP-AMPARs in mouse A2-ACs. In contrast to the inner nuclear layer, the ganglion cell layer showed little cobalt uptake in the *wt* or *rd10* retina (n=3 mice). We identified GCs using the pan-GC marker RBPMS, α-type GCs with neurofilament H (NFH), and the ON-sustained subset of α-GCs by their co-staining for NFH and calbindin (Fig. 7D-E). Although a sparse subset of GCs showed cobalt uptake, we saw no qualitative difference between *wt* and *rd10* retinas and no evidence for cobalt uptake in α-GCs. Since CP-AMPARs are not upregulated in GCs in the *rd10* retina, the functional effects of the CP-AMPAR blocker cannot be explained by direct effects on GCs.

**Figure 7:**
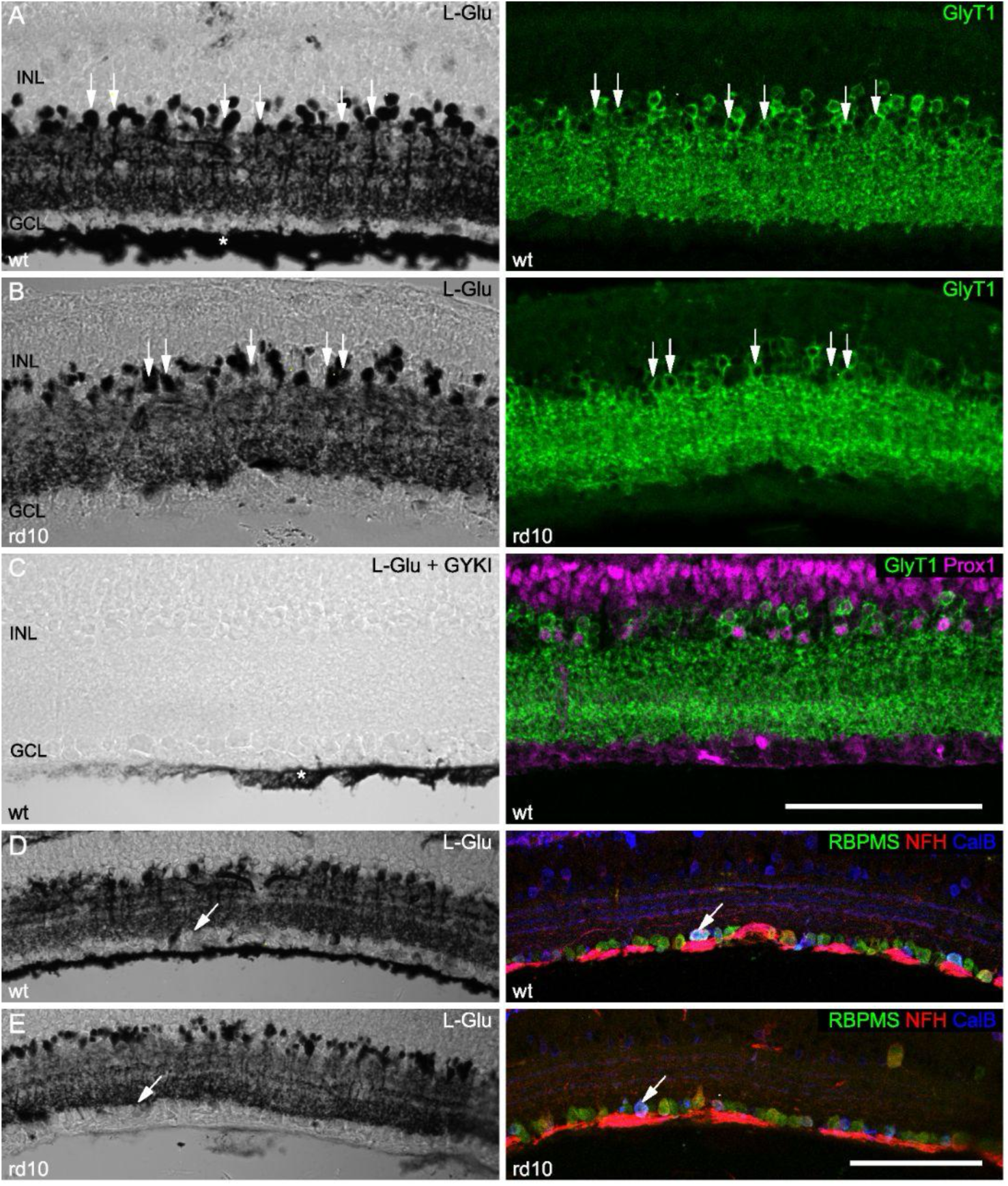
GCs lack cobalt uptake in *wt* or *rd10* retina. A,B. *Left panels.* L-Glu mediated cobalt uptake in GlyT1 positive amacrine cells (L-Glu) in *wt* (A) and *rd10* (B) retina. Arrows show examples of GlyT1+ ACs that show cobalt uptake, some of which have a thick proximal dendrite consistent with A2-AC morphology. **C.** Lack of L-Glu mediated cobalt uptake in the presence of the AMPA receptor antagonist, GYKI 53655. **D,E.** Lack of cobalt uptake in most GCs in the *wt* and *rd10* retina. Left panels show cobalt, right panels show immunostaining of the same retinal region. Arrows show examples of putative ON-sustained α GCs labeled with RBPMS, NFH and calbindin (CalB). Asterisks show non-specific cobalt staining associated with vitreous adherent to the retinal surface. INL, inner nuclear layer; GCL, ganglion cell layer; Scale bar in C applies to A-C, in E applies to D-E = 100 µm.

## Discussion

Aberrant GC activity emerges in a variety of rodent models of photoreceptor degeneration and has been attributed to changes in the inner retinal A2-AC network (Menzler and Zeck 2011; Choi et al. 2014; Biswas et al. 2014; Margolis et al. 2014, 2008; Borowska, Trenholm, and Awatramani 2011). Consistent with these prior studies, we find that aberrant activity in the *rd10* retina arises presynaptic to the GCs and further demonstrate that it is driven, at least in part, by excitatory input from RBCs to A2-ACs. We show that suppressing this excitatory input to A2-ACs improves the SNR of residual cone-driven signals in the *rd10* retina. We discuss the proposed mechanistic basis of these findings and potential implications for the treatment of retinal degeneration.

### Spontaneous activity is driven by CP-AMPAR mediated RBC inputs to A2-ACs

We found that A2-ACs in the old and young *rd10* retina were hyperpolarized by a CP-AMPAR blocker. Since most of the glutamatergic input to A2-ACs arises from RBCs, this result suggests that both RBCs and A2-ACs are tonically depolarized in the *rd10* retina. Consistent with this finding, RBCs are relatively depolarized in the *rd10* retina compared to wild type retina (Li et al. 2024). However, the impact of IEM on A2-AC oscillations was variable, causing a shift in oscillation frequency in some cells and complete suppression of oscillations in others. This result is consistent with prior studies in *rd1* mice showing that A2-AC oscillation frequency varies between 5-15 Hz (Choi et al. 2014; Yee, Toychiev, and Sagdullaev 2012; Trenholm et al. 2020) and depends critically on resting potential, with complete suppression of oscillations occurring only for the most hyperpolarized potentials (Choi et al. 2014). Our results suggest that tonic RBC input varies across individual A2-ACs in the degenerating retina (Fig. 1), presumably due to the uneven extent of photoreceptor degeneration. IEM also appeared to have more of an effect on A2-AC oscillations and On-α GC aberrant activity at the early compared to late *rd10* timepoints (Fig. 2), which may reflect a greater contribution from AMPA-receptor independent mechanisms, such as gap junctional input from On-CBCs to A2-ACs at later stages of degeneration (Toychiev et al. 2013). The localization of the CP-AMPA receptors to the A2-ACs precludes a direct effect of the IEM on the GCs (Fig. 6).

What drives RBC depolarization after rod loss? Since glutamate activation of mGluR6 receptors closes the downstream TRPM1 channel, the absence of glutamatergic input from photoreceptors could leave RBCs in a relatively depolarized state. However, rod degeneration leads to structural, molecular, and functional changes in RBCs including loss of synaptic mGluR6 receptor expression, dendritic remodeling, and formation of ectopic synapses with cones (Gargini et al. 2007; Puthussery et al. 2009). Although some extrasynaptic mGluR6 expression remains in RBCs after rod degeneration, mGluR6 receptor-mediated currents are markedly diminished, suggesting that most residual receptors are decoupled from the downstream TRPM1 cation channel (Puthussery et al. 2009). The loss of mGluR6 receptors could leave any membrane-associated TRPM1 channels constitutively open thus driving tonic RBC depolarization (Križaj et al. 2010). On the other hand, a recent study showed that low concentrations of the mGluR6 receptor agonist, L-AP4, improved the fidelity of GC signaling in the *rd10* retina, suggesting that some mislocalized mGluR6 receptors may remain coupled to TRPM1 (Li et al. 2024). In contrast to the mGluR6 receptors, somatic expression of TRPM1 persists after rod degeneration (Gayet-Primo and Puthussery 2015), although it is unclear whether these channels are in the plasma membrane or the endoplasmic reticulum pool (Agosto et al. 2018). Degeneration-induced changes in ion channel expression, such as the loss of BK channels (Schilardi and Kleinlogel 2021), could also contribute to RBC depolarization through a cell-intrinsic mechanism. Further experiments are needed to address these possibilities.

### Pathway-specific changes in GC activity during degeneration in rd10 mice

The results revealed two distinct pathway-selective changes during retinal degeneration. First, was an increase in the baseline spiking, which was only observed in the On-α GCs. The second was an unexpected increase in the amplitude of evoked responses, which again was only observed in the On-α-GCs.

As in prior studies, we found that spontaneous activity in *rd10* GCs increased with age (Stasheff, Shankar, and Andrews 2011). However, amongst the α-type GCs studied here, only the On-sustained cells showed a significant increase in spontaneous activity, with age. This age-dependent increase was seen in both the *wt* and *rd10* retinas, but the underlying mecchanisms are unclear. The selectivity for the On-pathway can be explained by the connectivity of A2-ACs. Tonic depolarization of the de-afferented rod-bipolar cells will produce tonic depolarization of A2-ACs (Fig. 2), which in turn will depolarize gap-junction coupled On-cone bipolar cells and bring On-GCs closer to spike threshold. In agreement, a prior study found that spontaneous firing increases in On-but not Off-GCs in P20 *rd10* retinas (Xiang et al. 2018). Similar to our results with IEM, (Xiang et al. 2018) showed that a low concentration of the competitive AMPA/KAR antagonist, CNQX, which will also block RBC input to A2-ACs, suppressed background firing in On-GCs. Such tonic depolarization is also evident in recent work showing that the threshold for activation of Off-GCs with light or electrical stimulation in *rd10* retina is higher than for On-GCs due to increased crossover glycinergic inhibition (Carleton and Oesch 2024; Dyszkant et al. 2024). It should be noted, however, that other studies have shown oscillatory or heightened spontaneous activity in Off-α GCs in the *rd1 and rd10* retina (Margolis et al. 2008, 2014; Ivanova et al. 2016). Our recordings in *rd10* retinas were limited to ages at which GCs retained light responsiveness. It is possible that additional mechanisms drive hyperactivity at later stages of degeneration. Indeed, *rd1* retinas degenerate much more rapidly than *rd10* and exhibit rhythmic oscillations in On- and Off-GCs that are less apparent in GCs of the *rd10* retina (Stasheff, Shankar, and Andrews 2011; Toychiev et al. 2013). The same A2-AC depolarization that drives baseline spiking in the On-pathway will hyperpolarize Off-CBCs through glycinergic “crossover inhibition”. It remains unclear why the crossover inhibition didn’t supress baseline firing in the Off-type cells.

The second pathway-specific effect was an unexpected increase in the amplitude of light-evoked On-responses in the *rd10* retinas with age, that was not seen in the Off-GCs, nor in any cells in the *wt* retina. The reasons for the differences are not clear, but suggest pathway-selective circuit-level compensation for the reduced photoreceptor input during degeneration (Care et al. 2020, 2019; Lee et al. 2021; Scalabrino et al. 2022; Leinonen et al. 2020; Ellis et al. 2023). The recent finding that voltage-gated calcium currents in type-6 On-CBCs decline during the progression of degeneration in both *rd1* and *rd10* mice (Ganzen et al. 2024) seems to be at odds with the potentiation of On-α GC responses seen here, suggesting that compensation may not be common to all On GC types.

### Site of action of CP-AMPAR antagonists in the retinal circuit

CP-AMPARs are expressed in horizontal cells (HCs) and A17 ACs (Osswald, Galan, and Bowie 2007; Chávez, Singer, and Diamond 2006; Percival et al. 2022) and can be upregulated in other neurons during ocular hypertension and glaucoma (Cueva Vargas et al. 2015; Xia, Nawy, and Carroll 2007; Wen et al. 2018; Sladek and Nawy 2020). The absence of CP-AMPARs on GCs, as suggested by the results of the cobalt assay (Fig. 6), rules them out as a site of action. Could the antagonist effects be due to blocking negative feedback from HCs to rod and cone photoreceptors (Barnes et al. 2020)? Light hyperpolarizes photoreceptors and suppresses glutamate release, thereby hyperpolarizing HCs, which is akin to the antagonist effect. Hyperpolarizing HCs will activate negative feedback, which will increase glutamate release from cones and hyperpolarize On-CBCs (and RBCs), but depolarize Off-CBCs. Thus, the CP-AMPAR antagonist should *decrease* tonic excitatory activity through the On-pathway but by the same token it should *increase* tonic excitation through the Off-pathway. However, this latter effect was not observed - the CP-AMPAR antagonist did not increase baseline firing of Off-GCs, indeed a slight *decrease* in baseline firing was seen in the old tOff-α cells, while the sOff-α cells showed no change. It is unlikely that these small, apparently anomalous effects on the Off-pathway originate in the outer layer, but we cannot rule out effects in the IPL circuitry.

HCs may also provide feed-forward GABAergic input that depolarizes On-CBCs and hyperpolarizes Off-CBCs owing to differences in somato-dendritic chloride gradients (Duebel et al. 2006). This site of action is unlikely because blocking GABA receptors did not recapitulate the effects of the CP-AMPAR antagonist. CP-AMPARs are also expressed at the RBC-A17-AC synapse where they mediate reciprocal inhibitory feedback onto the RBC terminal (Chávez, Singer, and Diamond 2006). This synapse is also an unlikely site of action because blocking CP-AMPARs will suppress reciprocal inhibition and in turn depolarize the RBC terminals, which would depolarize the A2-ACs, not hyperpolarize them as we have shown (Fig. 2). Taken together, the effects of blocking CP-AMPARs are inconsistent with an outer retinal origin and most consistent with suppressing transmission from the RBCs to A2-ACs.

### Comparison with other strategies to improve SNR in degenerating retinas

Our results show that low concentrations of a CP-AMPAR antagonist can improve the SNR of residual cone signals in On-α GCs. Consistent with our findings, application of low concentrations of the mGluR6 receptor agonist, L-AP4, also increased SNR in On-α GCs in the *rd10* retina (Li et al. 2024), presumably by suppressing tonic RBC excitation of A2-ACs. Since A2-ACs are coupled to most On-CBCs, albeit with a bias towards bipolar types 6, 7 and 5a (Tsukamoto and Omi 2017), CP-AMPAR blockers may have similar effects on other On-GC types, but this remains to be tested. Note that IEM appeared to have more marked effects on A2-AC oscillations and On-α GC aberrant spiking in young versus old *rd10* mice. Thus, from a therapeutic timing standpoint, the optimal window for treatment with such antagonists would most likely be in the period coincident with peak rod degeneration.

GC hyperactivity can be suppressed by pharmacological blockade or genetic knockout of gap junctions, leading to improved SNR of residual cone signals and improved optogenetic- or prosthetic-driven signals (Barrett, Degenaar, and Sernagor 2015; Eleftheriou et al. 2017; Toychiev et al. 2013). These effects are consistent with our findings and the hypothesis that RBCs drive aberrant activity through A2-ACs coupled to On-CBCs. However, gap-junctions are a less attractive therapeutic target than the CP-AMPARs because they contribute to a variety of physiological processes, such as discrimination of local vs global objects (Roy, Kumar, and Bloomfield 2017), motion sensitivity (Michael B. Manookin, Patterson, and Linehan 2018), orientation selectivity (Nath and Schwartz 2017), and rod-cone coupling (Raviola and Gilula 1973). Therefore, blocking gap-junctions could have extensive off-target effects on normal visual function. Moreover, gap-junctions blockers, such as meclofenamic acid, have anti-inflammatory properties and can alter voltage-gated potassium channel function (Peretz et al. 2007), further complicating their therapeutic utility. In comparison, we have shown that CP-AMPAR blockers would be expected to have limited and selective effects on retinal processing.

Our results align with prior work showing that hyperpolarizing A2-ACs can suppress aberrant activity in the context of retinal degeneration. For example, the M-type (Kv7, KCNQ) potassium channel activator, flupirtine, hyperpolarizes A2-ACs (Cembrowski et al. 2012), suppresses oscillatory activity in rd1 retina (Choi et al. 2014) and improves the signal-to-noise ratio of channelrhodopsin-driven signals (Barrett, Degenaar, and Sernagor 2015). The effects of flupirtine on native cone-driven signals has not been assessed in *rd* models, although persistence of rod-mediated signals was reported in the complexin 3 knockout mouse, which also shows relatively depolarized A2-ACs (Mortensen et al. 2016). Notably, KCNQ channel subunits are ubiquitously expressed in primate retinal ganglion cells as well as in a subset of amacrine cells (Peng et al., 2019, GEO: <u>GSE118480</u>, data not shown). Moreover, since flupirtine has reported activity on GABA_A_ receptors (Klinger et al. 2015, 2012), which are widely expressed in the inner retina, its therapeutic utility in the context of retinal degeneration remains to be empirically determined.

### Significance for human patients with IPRD

Our results suggest that CP-AMPAR antagonists may be useful therapeutic agents to improve visual processing in RP patients, even in the late stages of the disease when many patients retain some cone function (Jacobson et al. 2010; Gerth et al. 2007). Although direct evidence for aberrant GC activity in human RP patients is lacking, patients experience spontaneous photopsias, scintillations, and reductions in contrast sensitivity that may be linked to increased spontaneous GC activity (Stasheff 2018; McAnany et al. 2013; Bittner, Diener-West, and Dagnelie 2011). As in mouse models, cones degenerate slowly in the human RP retina leading to neuronal sprouting, and disorganization of the outer retina (B. W. Jones et al. 2011; Marc et al. 2007; B. W. Jones et al. 2016). Moreover, reductions in mGluR6 receptor expression are evident in human On bipolar cells after photoreceptor degeneration due to serous retinal detachment, and in a macaque model of acute photoreceptor loss (Gayet-Primo and Puthussery 2015; Patak et al. 2024). We recently showed that background and sustained spiking in On-midget and On-parasol GCs in the macaque retina is driven by the RBC → A2-AC circuit and can be blocked with CP-AMPA receptor antagonists (Percival et al. 2022). Our current findings indicate strong conservation of this retinal circuit from mouse to primate, suggesting the potential for translation to human therapies. Further work in primate models of photoreceptor degeneration will be valuable to test the efficacy of CP-AMPARs in improving GC signaling and behavioral performance (McGregor et al. 2022).

## Materials and Methods

### Animals and Tissue Preparation

All animal procedures were approved by the Institutional Animal Care and Use Committee of the University of California, Berkeley. We used homozygous C57BL/6J *wt* (JAX strain#:000664) and *rd10* (JAX strain#:004297) mice of either sex. Here, “young” animals are defined as postnatal (P) day P21-25 and “old” are P35-70. Animals were maintained on a 12-hour light/dark cycle and were dark-adapted for ∼2 hours prior to euthanasia. All subsequent procedures were performed under dim-red or infrared light (850 nm). Mice were deeply anesthetized with isoflurane and euthanized by cervical dislocation. Eyes were then immediately enucleated and the anterior eye and vitreous were removed under infrared illumination (850 nm) in Ames’ medium (U.S. Biologicals) equilibrated with carbogen (95% O_2_ and 5% CO_2_) at room temperature. For preparing retinal slices, the retina was hemisected and mounted on 0.45 µm nitrocellulose membrane with the ganglion cell side adherent to the paper.

### Single-Cell Electrophysiology

Single cell recordings were made on an Olympus BX51WI microscope with a 40x/0.8 N.A. water immersion objective under infrared illumination (850 nm), using a HEKA EPC10 amplifier. Cells were visualized with infrared illumination (870 nm) and Dodt gradient contrast optics using a 40x/0.8 N.A. or 20x/0.95 N.A. water immersion objective. Electrodes were pulled with borosilicate glass pipettes to a resistance of ∼10 MΩ. To record voltage responses in whole-cell current clamp, the internal solution contained (in mM) 128 K-methane-sulphonate, 9 KCl, 2 Mg-ATP, 1 Na-GTP, 1 EGTA, 10 Na_0.5_-HEPES and 2.5 Na_2_-phosphocreatine (pH 7.4). To record currents in whole-cell voltage clamp, the K^+^ was replaced with Cs^+^ to suppress voltage-gated K-currents. In most cases, the fluorescent dye Alexa Fluor 488 hydrazide (100 µM, Invitrogen) was included in the pipette to fill the cell and reveal its dendritic morphology.

Recordings from A2-AC were made in 300 µm thick vertical slices. Slices were made using a manual tissue chopper and mounted between grease tracks on a glass coverslip. Recordings were restricted to regions within ∼2 mm of the optic nerve head. A2-ACs were targeted based on their proximity to the inner nuclear layer, characteristic inverted pear shape of the somas, thick primary dendrite (Veruki and Hartveit 2002), and the presence of sodium spikelets during 10 mV depolarizing pulses.

Recordings from ganglion cells were made in hemisected, flat-mounted retina. Retinas were placed photoreceptor side down on a Whatman Anodisc (0.2 µm diameter; Cytiva) and held in place with a “harp” (Warner Instruments). Sustained and transient α-type retinal ganglion cells (GCs) were targeted based on their large soma size and characteristic spike responses (van Wyk, Wässle, and Taylor 2009). Loose cell-attached recordings were made with borosilicate glass microelectrodes (∼5 MΩ resistance) filled with Ames’ medium. Spike recordings were made with a HEKA EPC10 amplifier. In some experiments, cells were filled after spike recordings with an internal solution containing (in mM) 125 Cs-methane-sulphonate, 4 Na_2_-phosphocreatine, 10 CsCl, 2 Mg-ATP, 1 Na-GTP, 1 EGTA, 10 HEPES (pH 7.4) and 100 μM Alexa Fluor 488 to visualize cell morphology.

For pharmacology experiments, concentrated stocks of IEM-1460 (CP-AMPAR antagonist, Tocris Bioscience, 25 mM), GABAzine/SR95531 (GABA_A_ receptor antagonist, HelloBio, 50 mM); TPMPA (GABA_C_ receptor antagonist, Tocris Bioscience, 100 mM); strychnine (glycine receptor antagonist, Sigma, 10 mM) were prepared in ultrapure water and stored at −20°C. Unless noted otherwise, the final concentrations applied to the retina were 50 µM IEM-1460, 5 µM SR95531, 100 µM TPMPA, and 1 µM strychnine.

### Light Stimulation

A2-ACs in retinal slices were stimulated with diffuse light flashes from a light-emitting diode (LED) projected through a 40x/0.80 N.A. objective (Olympus). The peak emission wavelength of the LED was 505 nm, close to the peak absorption wavelength for rhodopsin in the mouse retina (500 nm). Stimulus intensity was modified by varying the duration of the LED flash between 10 µs and 5 ms to find the saturating flash intensity. The effects of IEM on the A2-AC light responses were tested at a single flash intensity, which delivered approximately 200 photons/µm^2^.

For GC recordings, light stimuli were generated using custom software written in Igor Pro 9.0 and the PsychoPy v3.0 toolbox. Stimuli were displayed on an LG LP097QX1 TFT-LCD retina display (2048×1536) at 60 Hz and projected onto the retina through a 20x/0.95 N.A. or 10x/0.30 N.A. objective (Olympus). The display intensity was linearized with a calibrated lookup table. Contrast was defined as (L_max_ - L_min_)/L_background_. Each cell was approximately centered on the stimulus screen by moving the soma to the predetermined center location. For contrast sensitivity experiments, full-field square-wave modulated stimuli (1 Hz) were presented over a range of contrasts (5-100%) on a gray background. Stimuli were presented in pseudo-randomized order. Each stimulus was repeated 5 times.

### Response measurements

EPSC amplitudes, estimated as the average amplitude of EPSCs from at least 5 stimulus trials, were obtained for each control and drug condition. EPSC amplitudes were measured as the difference between the baseline before the stimulus and the mean current during 10 ms segments centered on fixed time-points at the peak of the EPSCs.

After measuring the spike-times during each stimulus trial, peristimulus spike time histograms (PSTHs) were generated using a bin width of 20 ms. Baseline spike-rate was measured as the mean rate during 2 s immediately preceding the light stimulus. The peak spike rate during the light response was measured from the PSTH as the mean during a 30 ms segment centered on a fixed time point. The time point corresponded to the peak time of the response in the control condition. Signal-to-noise ratio (SNR) was calculated as (Peak-rate - baseline-rate)/s.d., where s.d. is the standard deviation of the baseline spike-rate calculated over 2 s immediately before the light stimulus.

Contrast-response functions were fit with the equation:

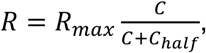

where *R* is the response amplitude, C is the contrast, *R_max_* is the maximum response amplitude and *C_half_* the half-maximal contrast. For many individual cells it was not possible to estimate *C_half_* from the equation above, because the responses did not clearly saturate within the contrast range. Therefore, in Fig. 5D we used an empirical half-maximal contrast, *C_1/2_*, which is the contrast at which responses reached 50% of the value at 100% contrast, as calculated from the fits of the equation to the data. These *C_1/2_* values will systematically underestimate the true values for large positive shifts in *C_half_*, however, the measurements correctly reflected changes in *C_half_*. Analysis was performed in Igor Pro 9 (Wavemetrics) using custom routines.

### Statistical analysis

Error bars/shading in the figures show ± 1 s.d. unless otherwise indicated. For statistical comparisons, datasets were first tested for normality using the Shapiro-Wilks test. Two-tailed paired or unpaired t-tests were used to compare datasets that conformed to a normal distribution. Welch’s correction was used for comparisons of datasets with unequal variances using the F-test of equality of variances. For non-normally distributed datasets, the Wilcoxon rank-sum or Wilcoxon signed-rank test was used as appropriate. In figures, significance was denoted by asterisks with ****, **, and * corresponding to p < 0.0001, 0.01, and 0.05 respectively. Not significant, n.s., p > 0.05.

We tested for significant changes in the amplitudes of the responses in the contrast-response relations using cluster-based permutation analysis as previously described (Maris & Oostenveld, 2007). At each contrast, control and drug amplitudes were compared using paired t-tests with *α* = 0.05 as the level of significance. Clusters were identified as 1 or more adjacent contrast points with p < 0.05. A cluster-statistic was obtained by summing the *p*-values for each cluster. We then generated a null-distribution by analyzing 5000 permutations in which control and drug data-points within each cell were randomly switched at each contrast. For each permutation we identified clusters, calculated the cluster-statistics, and added the maximum statistic to the distribution. The p values for observed clusters were calculated as the fraction of permuted cluster-statistics that were greater than or equal to the observed cluster-statistic, with p < 0.05 denoting significance.

### Immunohistochemistry

Immunohistochemistry was performed after cobalt assay and silver enhancement. Non-specific binding sites were blocked for 1 hr at 23°C in a buffer containing 10% normal donkey serum, 1% T×100, 0.025% NaN_3_ in PBS. Primary antibodies were diluted in 3% normal donkey serum, 1% T×100, 0.025% NaN_3_ in PBS and applied overnight at 23°C. Primary antibodies were RBPMS (1:500, Phosphosolutions Cat# 1832-RBPMS, RRID:AB_2492226); Neurofilament-H (1:2000, Covance Cat# SMI-32R-100, RRID:AB_509997); GLYT1 (1:4000; D.V. Pow, University of Queensland; Brisbane; Australia Cat# Glycine transporter 1 (GlyT1), RRID:AB_2314597). Whole IgG secondary antibodies, conjugated to Alexa Fluor 488 (Thermo Fisher Scientific Cat# A-21206, RRID:AB_2535792), Alexa Fluor 594 (Thermo Fisher Scientific Cat# A-21203, RRID:AB_2535789) or Alexa Fluor 647 (Jackson ImmunoResearch Labs Cat# 706-605-148, RRID:AB_2340476), were raised in donkey and used at a dilution of 1:800. Secondary antibodies were diluted in 3% normal donkey serum, 0.025% NaN_3_ in PBS and applied to tissue sections for 1 hr at 23°C. After final washes, samples were mounted in Mowiol. Confocal images were acquired on a Zeiss LSM 880 with a Plan-Apochromat 20x/0.8 air objective and the 488 nm, 561 nm and 633 nm laser lines and the silver-enhanced cobalt reaction product was imaged with the transmitted detector with Dodt contrast. Fluorescence channels were acquired sequentially to prevent cross-talk.

### Cobalt assay

CP-AMPAR expressing retinal neurons in *rd10* mice (4 mice, P48-P51) and *wt* mice (2 mice, P45-P55; 1 mouse, 8 months) were identified using the cobalt staining assay method described previously (Aurousseau, Osswald, and Bowie 2012). All steps were conducted in light-adapted retinas at 23°C and buffers were equilibrated with 95% O_2_ / 5 % CO_2_ for all steps before tissue fixation. Retinas were isolated as described above and stored in Ames’ medium for ∼1 hr. Hemisected retinas were first incubated for 50 minutes in an assay buffer containing (in mM) 57.5 NaCl, 5 KCl, 20 NaHCO_3_, 12 D-glucose, 139 sucrose, 0.75 CaCl_2_, 2 MgCl, and then transferred to assay buffer containing 5 mM CoCl_2_ plus either 10 mM L-glutamic acid (Sigma) or 10 mM L-glutamic acid + 80μM 1-(4-aminophenyl)-3-methylcarbamyl-4-methyl-3,4-dihydro-7,8-methylenedioxy-5H-2,3-benzodiazepine hydrochloride (GYKI 53655; Hello Bio). Retinas were then transferred to the assay buffer containing 2 mM EDTA (Invitrogen) to chelate excess extracellular cations. Next, Co^2+^ was precipitated with 0.24% ammonium sulfide (Sigma-Aldrich) in assay buffer. Retinas were washed in assay buffer and then fixed in 4% paraformaldehyde in 0.1 M phosphate buffer (PB) (Electron Microscopy Sciences 1578) for 90 minutes at 23°C. After washing in 0.1 PB, samples were cryoprotected in graded sucrose solutions (10-30%), embedded in Cryogel (Leica), sectioned at 16 μm, and stored at −20°C. The cobalt reaction product was visualized after 20-30 minutes of silver enhancement (Silver enhancer kit, Cat# SE100-1KT, Sigma-Aldrich).

